# Mutagenesis of two homologs ATP Binding cassette protein in tomato by CRISPR/Cas9 provide resistance against the plant parasite *Phelipanche aegyptiaca*

**DOI:** 10.1101/871087

**Authors:** Vinay Kumar Bari, Jackline Abu Nassar, Ayala Meir, Radi Aly

**Affiliations:** Department of Plant Pathology and Weed Research, NeweYa’ar Research Center, Agricultural Research Organization (ARO), Ramat Yishay, Israel

**Keywords:** CRISPR/Cas9genome editing, ATP Binding Cassette protein, strigolactone exporter, parasitic plant

## Abstract

Plant parasitic weed *Phelipanche aegyptiaca* and *Orobanche* spp. are obligate plant parasites that cause heavy damage to agricultural crop plants. Germination of the parasite seeds require exposure to specific chemical known as strigolactone [SL]. The plant hormone SL is derived from plant carotenoids via cleavage by *CCD7* and *CCD8* enzymes and exuded by the host roots to the rhizosphere. Here, we provide evidence that CRISPR/Cas9 mediated targeted mutagenesis of two homologues ATP Binding cassette (ABC) protein in tomato (*Solyc08g067610* and *Solyc08g067620*), significantly reduced the germination of the parasitic weed *P.aegyptiaca*. Constructs harboring specific single guide RNA were prepared and targeted against conserved region in the above tomato genes (*Solyc08g067610* and *Solyc08g067620*). Selected T0-mutated tomato plants showed different type of deletions at both locuses. Furthermore, genotype analysis of T1 plants showed that the introduced mutations stably inherited to next generation with no identified off-targets. Mutated tomato lines, showed more branching, enhanced growth of axillary buds, reduced length of primary stems and significantly reduced development of the parasitic weed *P. aegyptiaca* as compared to wild type plants. Moreover, in ^*ABC*^Cas9 mutants the expression level of the genes (*CCD8* and *MAX1*) related to SL biosynthesis were decreased, without alteration in the root extract orobanchol [SL] as compared to control plants. Development of genetic resistance based on genome editing of host key genes are novel approaches for enhancing host-parasite resistance. The current study offers insights into ABC protein homologs and mutagenesis of ABC protein that could be used in the development of efficient method to reduce parasitic weed germination.

## INTRODUCTION

The clustered regularly interspaced short palindromic repeats (CRISPR)/<u>C</u>RISPR <u>as</u>sociated protein <u>9</u> (CRISPR/<u>Cas9</u>) is a genome editing system direct precise target mutagenesis of specific gene and have significant applications in crop improvement^1^. The targeted gene knockouts mutagenesis achieved using CRISPR/Cas9, involved the double strand breaks (DSBs) induced by Cas9 are repaired by a non-homologous end-joining (NHEJ) which is a type of error-prone repair pathway that can introduce short deletions or insertions during DNA repair^2^. CRISPR/Cas9 system has three basic components: Cas9 endonuclease, CRISPR RNA (crRNA) and trans-activating crRNA (tracrRNA). tracrRNA:crRNA duplex specifically recognize the targeted complementary DNA strand and Cas9 endonuclease induce DNA break at a specific target site^3^. Recently both tracrRNA:crRNA were joined together to form a single guide RNA (sgRNA) containing an 18-20-nucleotide (nt) sequence which is selectively determines the target DNA sequence along with guide scaffold^4^. In addition, the 5’-NGG-3’protospacer adjacent motif (PAM) usuallyrequired for the CRISPR/Cas9 system^5^. The use of CRISPR/Cas9 system has been demonstrated to facilitate mutagenesis in diverse useful cropspecies, including Tobacco, sorghum, rice, cotton, arabidopsis, tomato, wheat^6–11^.

The obligate parasitic weeds (*Orobanche* and *Phelipanche* spp.) that completely depends on the host plant for nutritional requirement^14^, are major crop damaging agent in agricultural production^12,13^. The parasite attacks the host roots of several major economically important crops belongs to the Umbelliferae, Solanaceae, Compositae, Leguminosae and Cruciferae plant families^15,16^. The life cycles of *Orobanche* and *Phelipanchespp*. start with minute seeds that remain dormant and viable in the soil up to decades^17^. At early stage, the germination of the parasitic seeds required specific chemicals known as strigolactones (SLs) exuded by the host plant roots^18,19^. Preconditioning of parasitic seed with host exudate necessary for parasites seed germination. Once germination has been triggered by the specific chemicals^20^, the seed germinate and elongates towards the host root and develop a special connection known as haustorium^21^ that connects parasite directly to the vascular system of the host plant. The parasite grows into a bulbous structure known as tubercle and after 1 month of tubercle growth, a floral meristem is produced, which emerges above the ground to produce flower and disseminate seeds^22,23^.

SL, the germination inducer for parasitic weeds, derived from the carotenoid biosynthetic pathway, in which all-trans- β-carotene acts as the precursor for SL biosynthesis^24,25^. Several enzymes act sequentially in SL biosynthesis pathway: DWARF27 (*D27*) is a β-carotene isomerase; CAROTENOID CLEAVAGE DIOXYGENASE 7 (*CCD7*) and *CCD8* cleave 9-cis-β-carotene to produce the carotenoid precursor carlactone (CL). In tomato, MORE AXILLARY GROWTH 1 (*MAX1*) was identified as a class Ш cytochrome P450 enzyme that converts CL into carlactonoic acid (CLA), which is further converted in a species-specific way to either methyl carlactonoic acid (MeCLA)^26^, 4-deoxyorobanchol or orobanchol^27^. This core set of enzymes is mostly active in roots, and root-synthesized SLs are secreted into the rhizosphere or transported to the shoot^28^. Various types of SLs, e.g., strigol, 5-deoxystrigol, sorgolactone, solonacol, dideoxyorobanchol and orobanchol exuded from the host plant roots are known to stimulates germination of parasitic weeds seeds^29^. In previous study, reduced SL production in the root or exudation to the rhizosphere provide resistance to the host against parasitic weeds^30^. Previous studies with ABC-type transporters protein in *Petunia hybrida* reported a role of *PDR1* in the development of symbiotic arbuscular mycorrhizae and axillary branches, by functioning as a cellular SL exporter. Knockout mutants in *P. hybrid pdr1* were defective in SL exudation from their roots, resulting in reduced symbiotic interactions and parasitic growth of *P.ramosa*^31^. Here, we report the development of tomato plants which are resistant to the parasitic weed *P. aegyptiaca* using targeted mutagenesis of the two homologous ATP Binding cassette proteins (ABC) using CRISPR/Cas9 as a genome-editing tool.

## RESULTS

### Analysis of CRISPR/Cas9 mediated mutagenesis of two ABC homolog’s gene in tomato

To develop host resistance against root parasitic weeds, we chose to disrupt the *PDR1* orthologues in tomato plants using CRISPR/Cas9. To find the orthologous *PDR1*genes in tomato, we performed a protein BLAST search against tomato genome (Sol Genomics Network: https://solgenomics.net) using *Petunia hybrid PDR1*sequence. Similarity search outcomes showing maximum homology with the top two protein known as ABC subtype G (ABCG) transporters *Solyc08g067610*(ABCG44) (89.34%) and *Solyc08g067620*(ABCG45)(91.47%) which are highly conserved with respect to each other at protein level (86.00%). Recently, a genome-wide identification and expression analysis of genes encoding ABC proteins in tomato identified 154 genes putatively encoding ABC transporters and they suggests the physiological roles of ABC transporters ABCG44 and ABCG45 in SL transport which are highly expressed in root tissue^32^. Based on above conclusion, we chose to disrupt both genes using CRISPR/Cas9 system. To ensure successful editing of both genes, conserved sequence located in the twenty first exon of *Solyc08g067610* and *Solyc08g067620* gene were chosen, with a protospacer adjacent motif (PAM) sequence (NGG) at their 3 regions. A single ^*ABC*^sgRNA construct was designed using the selected sequence with PAM at their 3’ end and then combined into the Plant specific Cas9 vector (pHSE401)^33^ to target the *Solyc08g067610* and *Solyc08g067620* simultaneously (Fig. 1a and b). Agrobacterium mediated plant transformants were generated based on the presence of plant selectable marker hygromycin. A single independent T0 ^*ABC*^Cas9 transgenic plant were chosen which showed mutation at both the locus. To detect mutations induced by the Cas9 nuclease in the T0 plants, we perform the PCR at both the targeted site using genomic DNA from the T0 lines and sequenced. Sequence analysis of the amplified PCR products on the target sites of ABCG44 and ABCG45, showed multiple peaks, suggesting that they represent either heterozygous or biallelic mutants (Fig. 2a and b). Purified PCR fragments were also cloned into a TA cloning vector and again sequenced. Sanger sequence analysis revealed that mutation generated at ABCG44 locus are biallelic and at ABCG45 locus are heterozygous and contain three types of mutations: 8, 4 or 1nt. deletion. Interestingly, for ABCG44, both alleles lost wild type copy whereas ABCG45contains a single intact wild type copy after Cas9-mediated editing of the target (Fig. 2c-f). Since T0 transgenic tomato lines represent somatic mutation they were grown to maturity and self-pollinated to generate T1 progeny. Genomic DNA was extracted from the T1 progeny and the target region of Cas9 for both genes were amplified using PCR primers flanking the target sites. At least 30 plants from the T1 lines were examined for genotype at the target site using Sanger sequencing of target-site PCR products. In all T1 plants, mutations were conserved as shown in the T0 lines. Inheritance of the mutations by T1 plants, suggest that mutations resulted by genome editing activity were stably inherited in the next generation. The biallelic or heterozygous mutations were separated in T1 generation and some of the mutated lines were homozygous for mutation in both the target genes (ABCG44 and ABCG45). The biallelic mutation at ABCG44 locus were segregated in the T1 generation, and the ratios between the two mutations in a biallele were 1:2:1 12(8bp deletion):6(4bp deletion):10(biallelic). Similarly, the heterozygous mutation at ABCG45 locus were 1:2:1 12wt:6(1bp deletion):7(heterozygous) (Table S1). Moreover, examination of transgenic region in some of the T1 generation plants suggest that 48% (12/25) of T1 plants were detected to be transgene free and did not contains any foreign DNA sequence (Fig. S1 and Table S1). The putative off-target sites associated with ^*ABC*^sgRNA were evaluated by CRISPR-P program^34^. We analyzed three potential off-targets sites with high scores, in the T1 generations of Cas9-edited tomato plants. Sequencing of PCR products from these regions revealed no changes in the putative off-target sites in the Cas9-mutated plants (Fig. S2 and Table S2).

**Figure 1.**
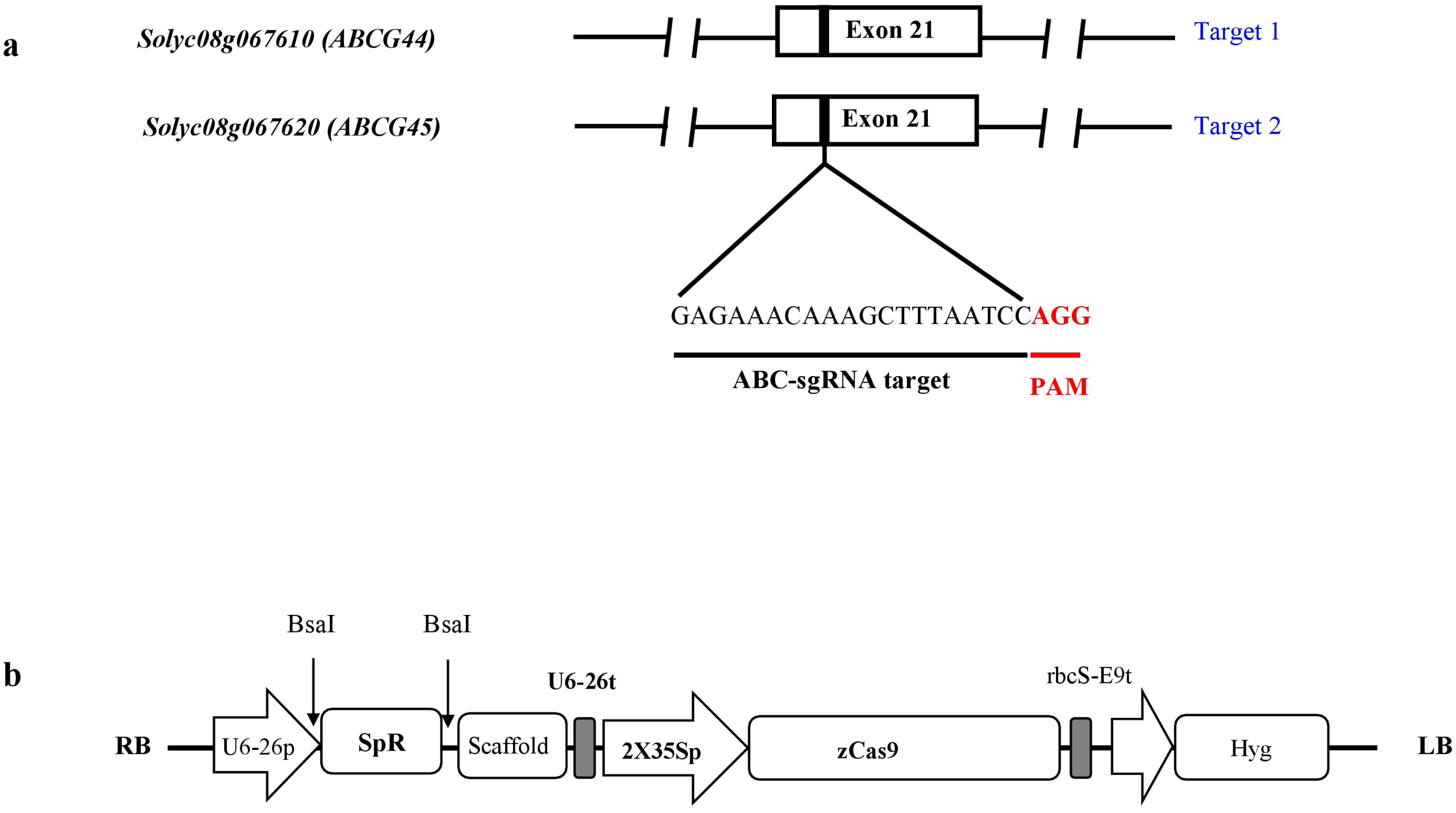
Schematic representation of the ^*ABC*^sgRNA target site and CRISPR/Cas9 vector construct used to mutate two ABC homologs gene. **(a)** Schematic representation of the tomato *Solyc08g067610* and *Solyc08g067620* genomic map and location of the ^*ABC*^sgRNA target site. The target sequence of ^*ABC*^sgRNA is shown in black color present in the 21^th^ exon; the PAM is shown in red. **(b)** The strong constitutive 2X35S promoter was used to express *Zea mays* codon-optimized Cas9 and the *Arabidopsis* U6-26 promoter was used to express ^*ABC*^sgRNA. sgRNA was cloned under *Bsa*I target site.

**Figure 2.**
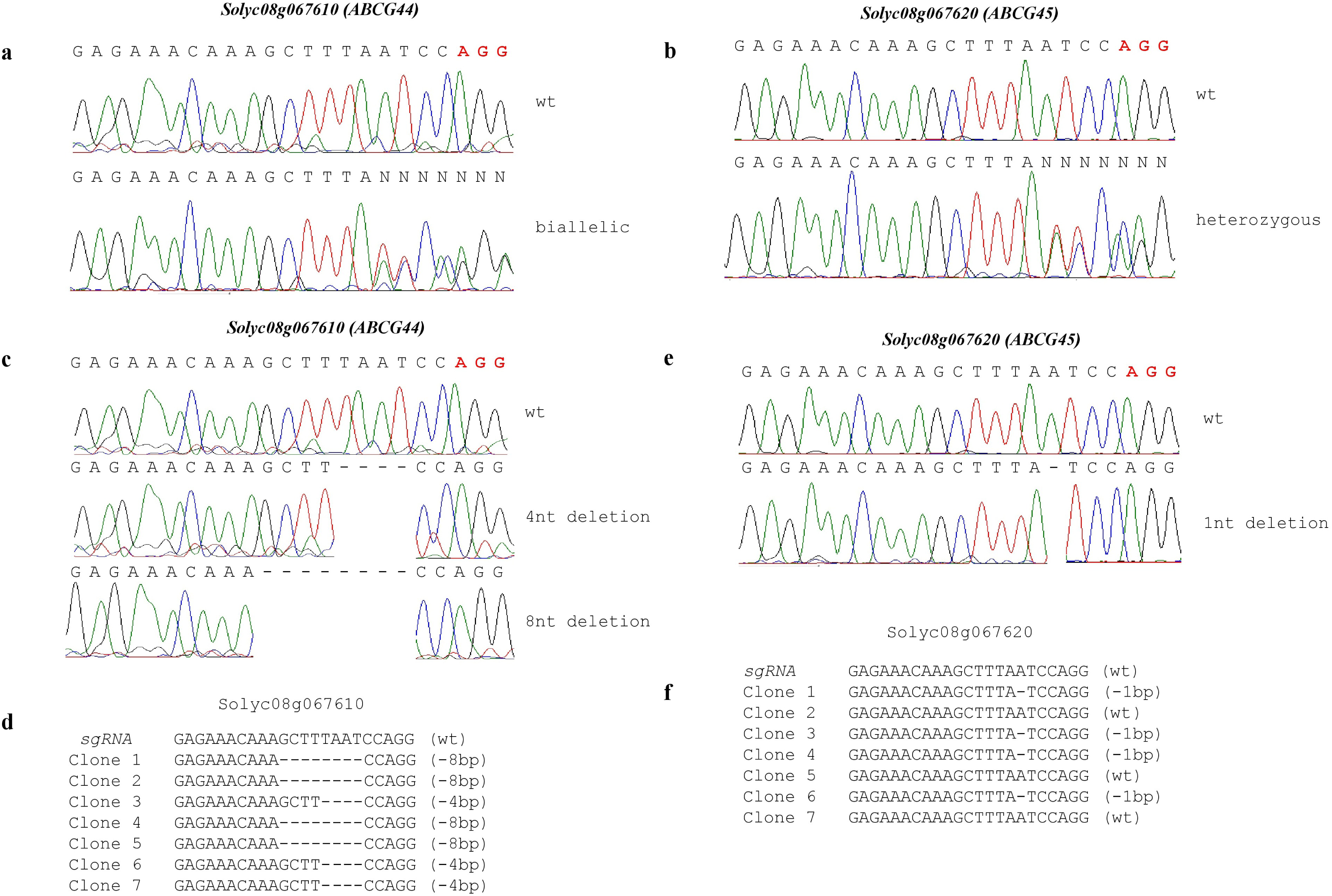
CRISPR/Cas9 generated mutation detection in T0-generation. **(a)** PCR fragments sequencing of *Solyc08g067610*targeted region of the T0 lines shown multiple peak as compared with the wild-type genome sequences (WT) **(b)** PCR fragments sequencing of *Solyc08g067620* targeted region of the T0 lines shown multiple peak as compared with the wild-type genome sequences (WT) **(c-d)** *Solyc08g067610* target regions were sequenced after PCR product cloned into the TA vector. Sanger sequencing of positive clones aligned with wild-type sequence, and type of mutation (indels) detected presented on the right side of the sequences. **(e-f)** *Solyc08g067620* target regions were sequenced after PCR product cloned into the TA vector. Sanger sequencing of positive clones aligned with wild-type sequence, and type of mutation (indels) detected presented on the right side of the sequences.

### Morphological phenotype *of^ABC^Cas9* mutated tomato plants

The plant hormones SLs regulate plant shoot architecture by inhibiting the outgrowth of axillary buds^35^. Despite their importance, it is not known how SL is transported. Previous studies with *Petunia hybrida* ABC protein *PDR1* reported that above ground, *pdr1* loss-of-function mutants excreted less SL in the rhizosphere have an enhanced branching phenotype, due to impaired SL allocation^31^. In addition, *NtPDR6*, a SL-transport deficient mutant generated through CRISPR/Cas9 in tobacco is known to exhibit increase in shoot branching, lateral roots and overall dwarfing^36^. Similar to previous reports, we also observed similar morphology in both the ^*ABC*^Cas9 mutated plants such as decreased shoot heights, branched shoots were significantly increased and more growth of axillary buds as compared to the wild type plants; however the mutagenic plant does not have significant differences in their root mass as compared to control plants (Fig. 3a-e and Fig. S3).

**Figure 3.**
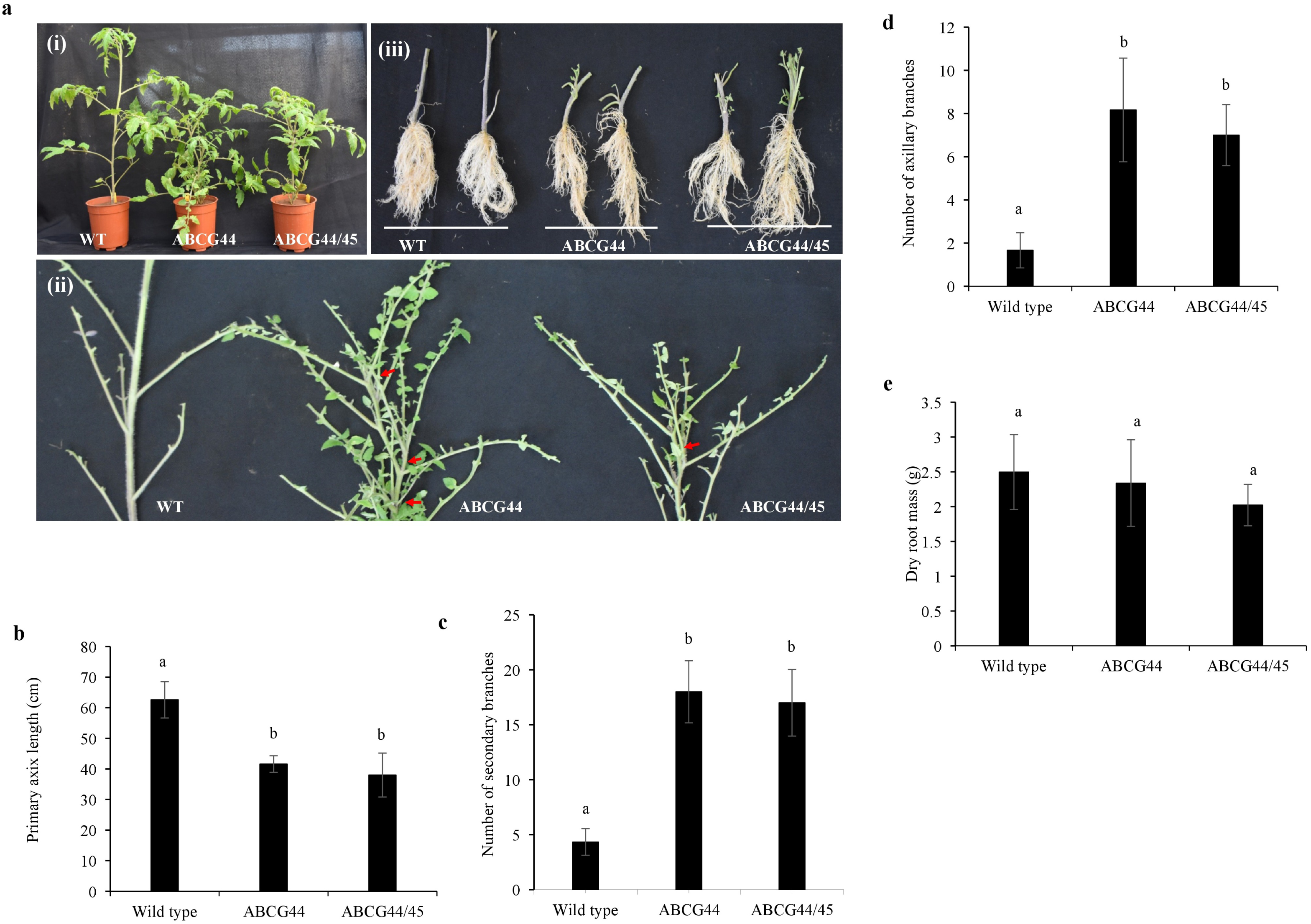
Morphological phenotypes of ABC knock out mutations induced by CRISPR/Cas9. **(a-i):** Normal vegetative growth of wild type (WT); *Solyc08g067620*-Cas9 edited T1 single mutant (ABCG44); *Solyc08g067610* and *Solyc08g067620* Cas9 edited T1 double mutant (ABCG44/45) plants. **(a-ii):** Difference between branching and axillary buds growth of the wild type and ^*ABC*^Cas9 mutated single mutant (ABCG44) and double mutant (ABCG44/45) plants **(a-iii):** Root morphology. **(b):** plant height after 2 month of growth average ±SE (n=8) **(c)**: Quantitative estimate of secondary branches in 2 month old ^*ABC*^Cas9 mutated tomato plant, average ±SE (n=5) **(d):** Quantitative estimate of axillary branches in 2 month old ^*ABC*^Cas9 mutated tomato plant, average ±SE (n=5) and **(e):** Quantitative estimate of dry root mass average ±SE (n=5), in ^*ABC*^Cas9 mutated and wild type plant. The different small letters not connected by same letter on each bar indicate statistically significant differences compared to wild-type plants (p-value < 0.05; Student’s t-test).

### Resistance of ^*ABC*^Cas9 mutated tomato plants to *P. aegyptiaca* and their orobanchol content

To analyze whether the ^*ABC*^CRISPR/Cas9-mutated lines confer resistance to *P.aegyptiaca*, selected mutated tomato lines from T1 generation showing the knockout phenotypes characteristics, were infested with *P. aegyptiaca* seeds. Homozygous for single or double mutation of T1 plants, were grown into small pots containing soil with *P. aegyptiaca* seeds and grown in a greenhouse with control conditions. Two separate experiments with five replicates per treatment were conducted. To measure the resistance of the mutated lines, we counted only fresh and viable parasite tubercles which are larger than 2 mm in diameter. Our results indicate that the numbers of attached parasite tubercles and shoots were significantly reduced in the ^*ABC*^Cas9 mutated lines as compared to the wild-type plants (Fig.4a). To further explore the relation between SL exudation by ABC protein in the ^*ABC*^Cas9 mutants and their resistance to *P. aegyptiaca* infection, we analyzed the total orobanchol content in the roots of wild-type and ^*ABC*^Cas9 mutated plants. We did not find any significant differences between Orobanchol content in the single or double ^*ABC*^Cas9 mutated plants as compare to the wild type (Fig. 4b).

**Figure 4.**
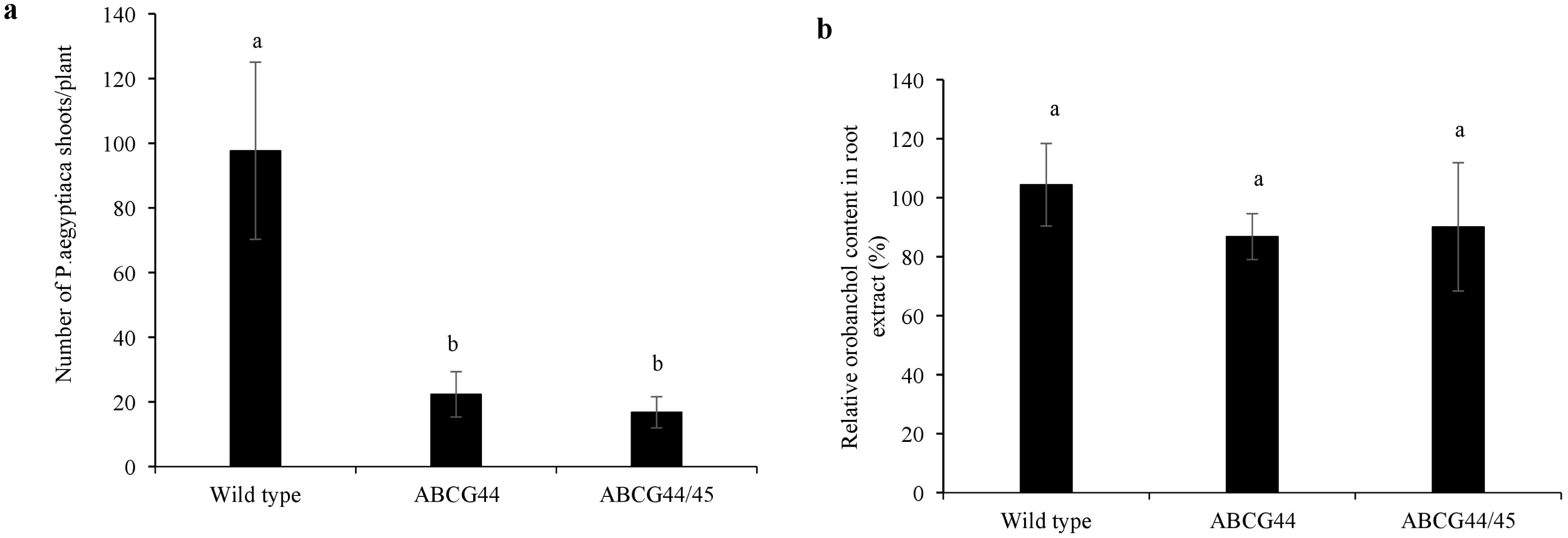
Resistance of tomato against *P.aegyptiaca* and Orobanchol content in the ^*ABC*^Cas9 mutant plant roots. To evaluate host resistance to the parasite, host roots of the tomato wild-type (WT) and ^*ABC*^Cas9 mutated T1 plants were washed after 3 months of infection with *P. aegyptiaca*. Tubercles larger than 2 mm in diameter were considered for analysis. **(a)** Number of *P. aegyptiaca* tubercles and shoots attached to the mutant and non-mutant tomato plants in the pot assay. Bars represent average of two experiments with five independent plants of each group ± SD values. **(b)** Orobanchol contents in the roots extract of tomato ^*ABC*^Cas9 edited single (ABCG44) and double (ABCG44/45) mutant plant as compared to wild type. LC-MS/MS analysis was done with three independent biological samples from each mutant. Data presented as average two independent experiments±SD.

### ^*ABC*^Cas9 mutation altered the expression of *CCD8* and *MAX1* genes involved in SL biosynthetic pathway

Carotenoid biosynthetic pathway derivatives all trans-β-carotene act as precursor for production of SL and this pathway are under negative feedback control by SL content^37,38^. Our Cas9 mutants are defective in SL export which can cause retention of SL in the host roots which might affect feedback regulation of SL pathway. Hence, we are interested to discover whether defective SL export affects SL biosynthetic and upstream carotenoid pathway. A simplified diagram showing possible routes for SL transport in the root, location of ABC transport protein and SL biosynthesis pathways was illustrated (Fig. 5a and b). Since *CCD8* and *MAX1* catalyze a key step in SL biosynthesis; we to analyzed the expression of *CCD8* and *MAX1* in ^*ABC*^Cas9 mutants. Interestingly, *CCD8* and *MAX1* showed decreased expression in ABCG44 and ABCG44/45 ^*ABC*^Cas9 in edited T1 plant as compared to the wild type (Fig. 6a) suggesting that, ^*ABC*^Cas9 mutation affect expression of SL biosynthetic genes. This data also provide a basis to explain the highly branched shoot and enhanced growth of axillary buds in ^*ABC*^Cas9 mutated plants. To explore whether ^*ABC*^Cas9 mutation also affect the carotenoid content in the roots, the content and type of carotenoid present in wild type and ^*ABC*^Cas9 mutated lines were analyzed. Interestingly, no significant difference in total carotenoids and its derivative was observed in ABCG44/45^*ABC*^Cas9 double mutant plants; however ABCG44 ^*ABC*^Cas9 single mutant plants shown increase in total carotenoids including Lutein and β-carotene (Fig. 6b). To further confirm the above results, we analyzed the expression of prominent gene Phytoene desaturase-1 (*Solyc03g123760*) and Lycopene cyclase-β (LCY-β) involved in the carotenoid. No significant change in expression of *PDS1* and *LCY-β* was observed in both mutants; demonstrate that altered carotenoid profile in the root of ^*ABC*^Cas9 single mutants does not involved alteration in gene expression in *PDS1* or *LCY-β*. In conclusion, we generated SL export defective mutants in tomato, which are resistance against parasitic weeds *P. aegyptiaca* due to inability to export SL to the rhizosphere.

**Figure 5.**
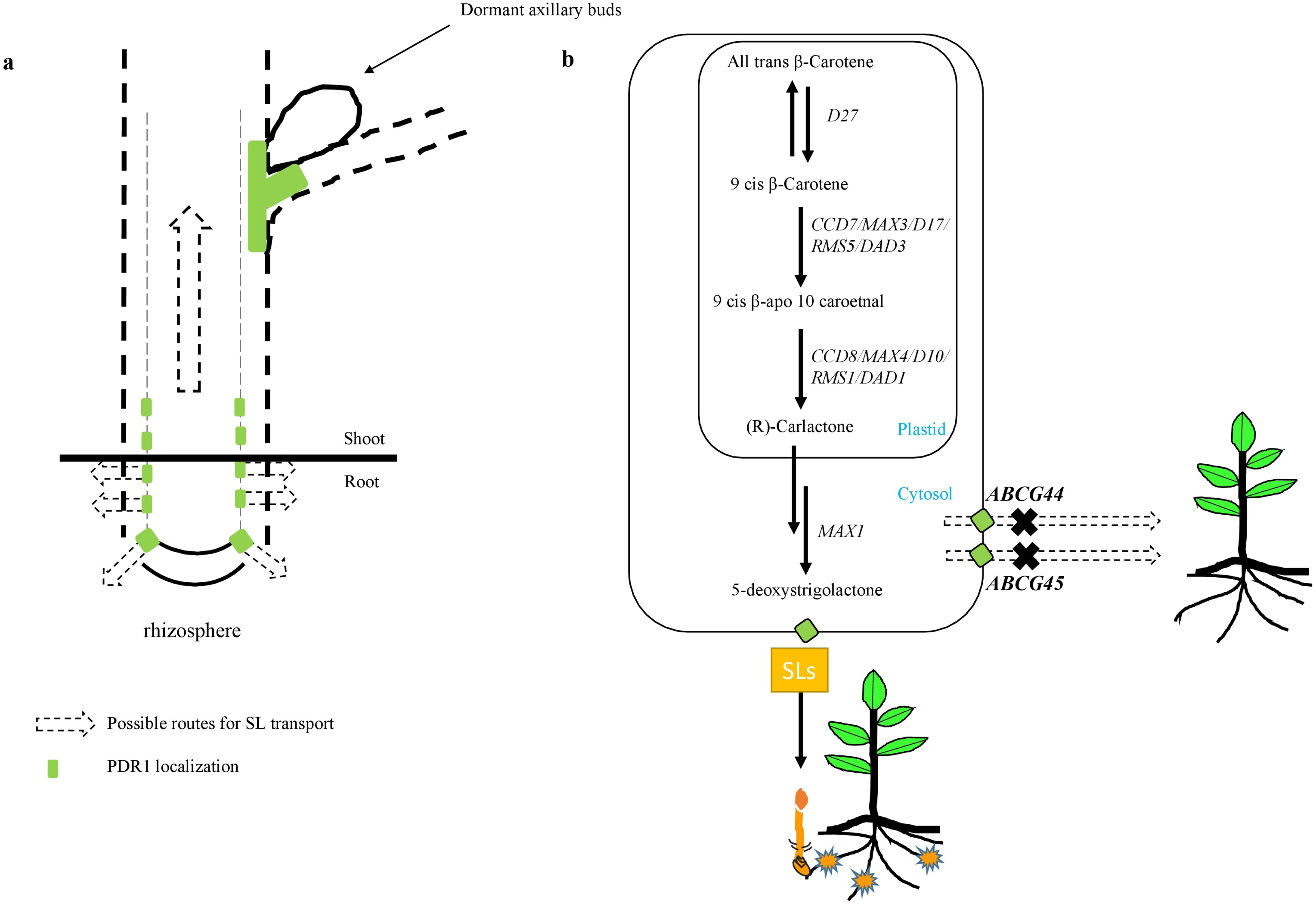
Diagram showing root tissue and the localization of ABC protein, SL biosynthesis pathway inside cell and export by ABC protein. **(a)** Diagram showing details of root tissue, possible SL transport pathways and localization of ABC protein (ABCG44 and ABCG45) in host root tissue. **(b)** SL biosynthesis and export protein shown inside a root cell. The enzyme belongs to SL biosynthesis are *D27*, carotenoid cleavage dioxygenases 7, 8 and *MAX1; ABCG44* and *ABCG45* are ABC protein homologs involved in SL exudation which leads to germination of *Phelipanche aegyptiaca* seeds.

**Figure 6.**
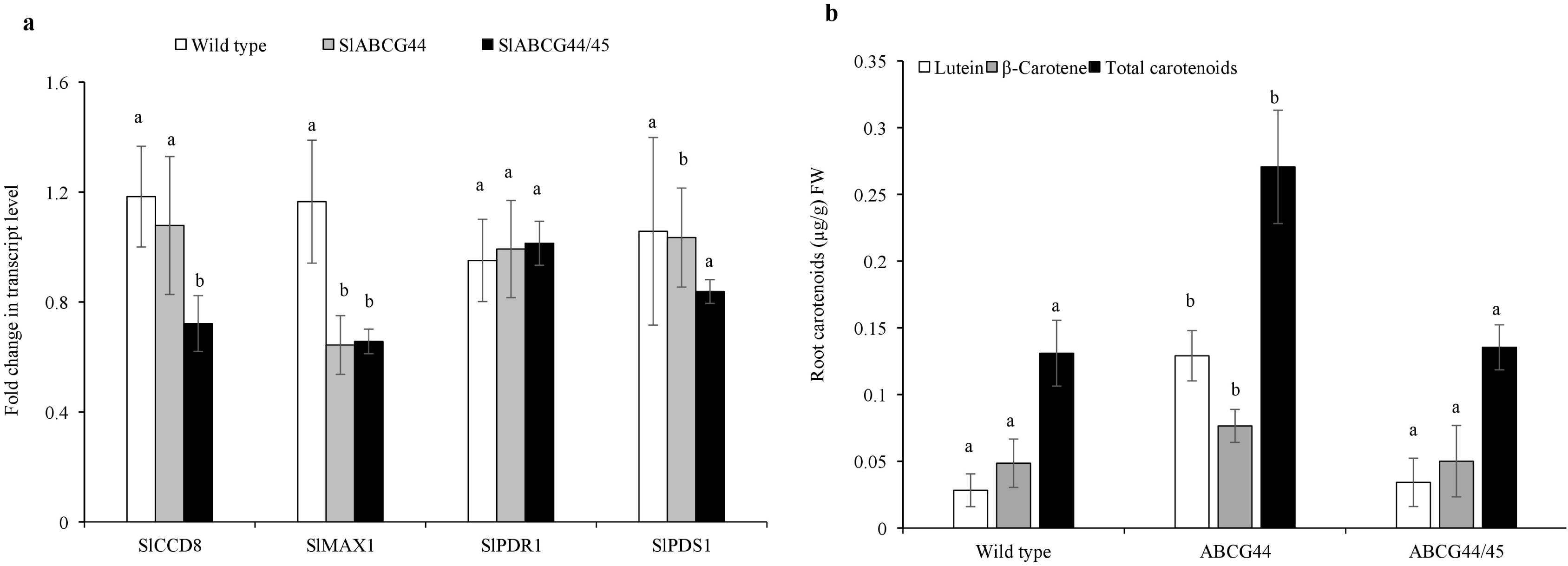
Carotenoid contents and expression analysis of selected genes involved in SL biosynthesis in ^*ABC*^Cas9 mutant plants. **(a)** Quantitative analysis of transcript levels in roots of ^*ABC*^Cas9 edited mutant and wild type (WT) plants using quantitative real-time PCR. Expression level of transcript was displayed after normalization with internal control tomato elongation factor-1α (EF1-α). Bars with different letters are significantly different from each other (student t-test at p< 0.05). Results represent the average of at least three independent experiments with three technical repeat ± SE (n=3), and expressed as fold increase in expression of transcripts relative to the control plants. **(b)** Carotenoid content in tomato roots of control and^*ABC*^Cas9 mutant plants. Values are based on the analysis of 2 months old plants grown in green house, bars represent the average of two experiments, was done with the pooled sample from three independent plant roots for each mutants ±SE. Statistical differences between the wild-type control were calculated with Student’s t-test (P<0.05). Different small letter on bar indicate a significant difference between the ^*ABC*^Cas9 edited mutant and the WT plants.

## DISCUSSION

The parasitic weeds (*Phelipanche* and *Orobanche* spp.) are concealed underground and exerts the greatest damage prior to its emergence; therefore, the majority of field loss may occur before diagnosis of infection. The lack of effective control strategies for parasitic weeds leaves farmers without viable options for protecting their crops. Therefore, development of parasite-resistant crops required to avoid parasitic weed growth. We hypothesize that mutagenesis of two homologues ATP Binding protein in tomato (*Solyc08g067610* and *Solyc08g067620*), could be used in the development of efficient method to reduce parasitic weed germination.

Previous work with ATP Binding cassette protein reported existence of SL transporter *PDR1* in *Petunia hybrida*, a closest homologue of Arabidopsis ABCG40 (also known as PDR12) which transports abscisic acid^42^, however, in contrast to Arabidopsis ABCG40, *P. hybrida PDR1* is not regulated by abscisic acid^31^. In subsequent study they reported that the *PDR1* exhibits asymmetrical localization in petunia root tips and *PaPDR1* mutants displayed increased shoot branching, altered orobanchol content in the root exudate and significantly decreased germination rate of *Orobanche ramosa*^31^, whereas the SL levels in root extracts were not affected. Thus, indicating that *Petunia pdr1* mutants are not defective in SL biosynthesis and *PDR1* functions as SL export carrier^43^. Our results were consistent with the previous studies; we observed similar morphological phenotype in the ^*ABC*^Cas9 tomato lines. Mutated-plants displayed decrease in plant height, increased shoot branching and more growth of axillary buds. Our previous observations with *^CCD8^Cas9* mutants suggest that^CCD8^Cas9 mutants are more branched than ^ABC^Cas9 mutants generated in the current study. More bud outgrowth was observed in ^ABC^Cas9 mutants than in the wild-type plants, causing longer branches at node (three to five). This early and vigorous bud outgrowth was also observed in pdr1-RNAi lines^31^. In contrast to ^ABC^Cas9 mutants, *dad1* mutants shown full branches from all nodes with retarded growth compared to the wild type. The integration of auxin and SL signaling seems to be involved in this phenomenon^44^. The mild branching phenotype of ^ABC^Cas9 mutants as compare to the SL-biosynthesis mutant *ccd8* suggests that residual transport and/or locally produced SL or other SL derivatives possibly compensate for defective SL transport in the shoot.

To determine host resistance to *P. aegyptiaca*, we used a pot system containing soil infested in advance with *P. aegyptiaca* seeds. We showed a significant decrease in the number of parasite tubercles and shoots growing on mutated tomato lines relative to the control treatment (the non-mutated wild-type plants were highly susceptible to the parasite infestation).

To date, several unique SLs, including orobanchol and solanacol, have been reported in the root exudates and extracts of tomato^45,46^. However, orobanchol is a major SL in tomato root exudates^46^ and is a specific germination stimulant for *P. aegyptiaca*. We determined orobanchol content in the roots of ^*ABC*^Cas9 mutated lines. Orobanchol levels in the ^*ABC*^Cas9 mutated plant relative to the wild type was similar. In addition to the previous studies, here we discovered that ^*ABC*^Cas9 mutants affected expression profile of some candidate genes involved in SL biosynthesis therefore, carotenoid biosynthesis was altered. Previous studies demonstrated that genes involved in SL-biosynthesis pathways are regulated by feedback inhibition. Therefore, decrease in SL content might affect the transcript levels of *DAD1* of Petunia, *D10* of Rice or *RMS1* of Pea in the respective mutant background^37,38^. Based on these results, we expect that ^*ABC*^Cas9 mutants will affect the expression of SL biosynthetic gene (*CCD8, MAX1*) transcripts. To measure the change in gene expression, we analyzed its relative transcript level using quantitative real-time PCR. Interestingly, the transcript level of *CCD8* and *MAX1*, both was substantially decreased in the single (ABCG44) or double mutated (ABCG44/45) tomato lines relative to the wild-type plants. We hypothesize that following mutation in SL export gene using CRISPR system resulted in absence of SL exudation that will restore feedback regulation due to retention of SL in the host roots. Recent study utilized CRISPR/Cas9 to engineer the rice plant architecture through mutagenesis of rice *CCD7* gene, showed reduced germination of the parasitic weed *Striga hermonthica*^47^. Similarly, In our previous studies, we have showed that CRISPR/Cas9 mediated gene editing of *CCD8* in tomato facilitates suppression of the plant parasite *P.aegyptiaca* growth^48^. In this study, CRISPR/Cas9 was used to precisely knock out the two *ABC* gene homologs using one single sgRNA in tomato. Parasitic weed resistance based on the editing of specific target genes using CRISPR/Cas9, can be easily applied to other susceptible crops and this method could be effective with other host plant, to develop parasite resistance. In addition, the host plants are not considered genetically modified organisms due to lack of any foreign DNA sequences in the mutated host plants^49^. One of the major challenges associated with CRISPR/Cas9 system is off-target cleavage due to non-specificity of the selected sgRNA with other few base pair mismatched target site within the genome^50^. Thus, this might affect the function of other non-specifically targeted gene in combination to the selected gene and may result in genomic instability. Although, to avoid off target effects several modifications of the Cas9 and advanced sgRNA designing methods have been developed which is still considered as major limiting factor with CRISPR/Cas9 technology^51,52^. In conclusion, we demonstrate that CRISPR/Cas9 mediated mutagenesis of two ABC homologs gene, reported to be involved in SL-exudation from the roots, provide genetic resistance to the parasitic weed *P. aegyptiaca* in tomato.

## EXPERIMENTAL PROCEDURES

### Materials and growth conditions

Tomato cultivar M182 was selected for transgenic plants construction. Sterile seedlings grown on 1/2 Murashige and Skoog (1/2 MS) medium with 0.7% phytagel, and they were placed on tissue culture chambers with temperature of 23 °C (day/night), light cycle of 16/8 h (day/night),. After ten days, cotyledons with 1 to 2-mm long petioles were cut from both side and used as explants in Agrobacterium-mediated transformation.

### ^*ABC*^sgRNA design and CRISPR vector construction

The tomato ABC protein *Solyc08g067620* and *Solyc08g067610* was chosen as the target for Cas9 editing. For developing the CRISPR/Cas9 gene editing constructs, a common sequence site was selected in the twenty one exon of both gene*Solyc08g067620* and *Solyc08g067610*. Two complementary oligos with 4bp overhangs (TmABC-sgRNA-F and TmABC-sgRNA-R were specifically designed manually and used for Cas9 targeted editing (Table S3). The target sequences for *Solyc08g067620* and *Solyc08g067610* are completely identical, and the GC content of sgRNA was 35%. The oligo pairs were first annealed to produce a double-stranded fragment with 4-nt overhangs at both ends and then ligated into the *BsaI* digested pHSE401, a CRISPR/Cas9 vector with maize codon optimized Cas9 under 35S promoter with hygromycin as plant selectable marker. To verify correct sgRNA inserted vector, independent transformants were PCR amplified and sequenced using HSE401-F and HSE401-R oligos. The construct was transformed into tomato cultivar M182 using *Agrobacterium tumefaciens* strain EHA105.

### Agrobacterium-mediated transformation of tomato plants

The transformation of tomato was conducted as previously described, with modifications^53^. The cotyledons were pre-cultured for 2 day in dark with Murashige and Skoog (MS) medium containing 100μM Acetosyringone, 0.1mg/L IAA, 1mg/L Zeatin and 0.7% phytagel subsequently infected with the Agrobacterium strain EHA105 (optical density at 600 nm less than 0.5) by immersion for 20 min. The explants were co-cultivated with Agrobacterium strain EHA105 for three days in dark and then transferred to the MS medium supplemented with 0.1mg/L IAA, 1mg/L Zeatin, 10μg/ml hygromycin, 300μg/ml timentin and 0.7% phytagel for a week. Hygromycin-resistant shoots regenerated were transferred to tissue culture bottles containing the subculture medium with 0.1mg/L Zeatin. Developed vegetarians were transferred to the half strength MS rooting substrate with the addition of 2mg/L IBA. After three months, hygromycin-resistant plantlets were obtained and used for subsequent analysis.

### Mutant detection and genotyping

Tomato leaves and roots genomic DNA was extracted using a Bio-Basic plant genomic DNA extraction kit and the genomic DNA flanks containing the target sites were amplified using the specific primers ABC610-Int-F &ABC610-Int-R or ABC620-Int-F &ABC620-Int-R and then run on a 2.0% agarose gel using electrophoresis. Image was acquired using DNR MiniLumi with UV light system. For genotyping the plants, PCR products amplified with the above primers were directly sequenced using appropriate primer. For PCR product cloning, pGEM^®^-T kit from Promega were used. The primers used in this study, are presented in Table S3.

### Analysis of off-target mutations

The potential off-target sites associated with ^*ABC*^sgRNA target sequence were analyzed with the CRISPR-P program^34^. Three sites with the highest off-target probability score were selected for the target. The genomic region flanks upstream and downstream to the off-target sites (300-400bp) was amplified and sequenced with specific primers using Sanger sequencing.

### RNA isolation and quantitative real-time PCR

Total RNA from tomato roots was extracted using spectrum plant total RNA kit (Sigma-STRN50-1KT) according to the manufacturer’s protocol. 500ng of total RNA was used to obtained cDNA according to the protocol of Quanta Bioscience cDNA Synthesis Kit. Quantitative real-time PCR (qRT-PCR) was performed in a volume of 10μl using PerfeCTa^®^ SYBR^®^ Green FastMix^®^, ROX™ (Quanta biosciences) with 5 times diluted cDNA as template. Tomato elongation factor 1-α was used as an internal reference gene. Specificity of the primers was confirmed by melting curve analysis. The generated Ct values of target genes were normalized to the Ct value of housekeeping EF1-α gene. Relative expression was calculated using 2^−ΔΔCt^ method and expressed as fold increase with respect to control^54^.

### Statistical analysis

JMP Pro 14 software and Microsoft Excel 2013 were used to perform standard statistical tests. All the experiments were repeated independently for at least three times. Error bars represent standard deviation from the mean of biological triplicate samples. Significant differences were determined by student t-test at p ≤ 0.05 (small letters) and represented as small letter on the bar.

## AUTHER CONTRIBUTIONS

V.K.B. and R.A. together conceived and planned the study. V.K.B. performed the target design, vector construction, transgenic plant generation, mutational and qRT-PCR analysis. J.A.N. performed *P.aegyptiaca* infestation analysis. V.K.B and A.M. performed the carotenoid analysis. V.K.B. and R.A. finalized the data and wrote the manuscript.

## ACKNOWLEDGMENTS

This research work was supported by research grant no. IS-4622-13 from BARD, the United States – Israel Binational Agricultural Research and Development Fund. Additional support was by research grant no. 132-1919-18, received from Chief Scientist of the Ministry of Agriculture and Rural Development (Israel). V.K.B. is grateful to the ARO-Volcani Center, Agricultural Ministry of Israel for providing the Postdoctoral fellowship.

## Supplementary Information

**Supplementary Fig. S1.**
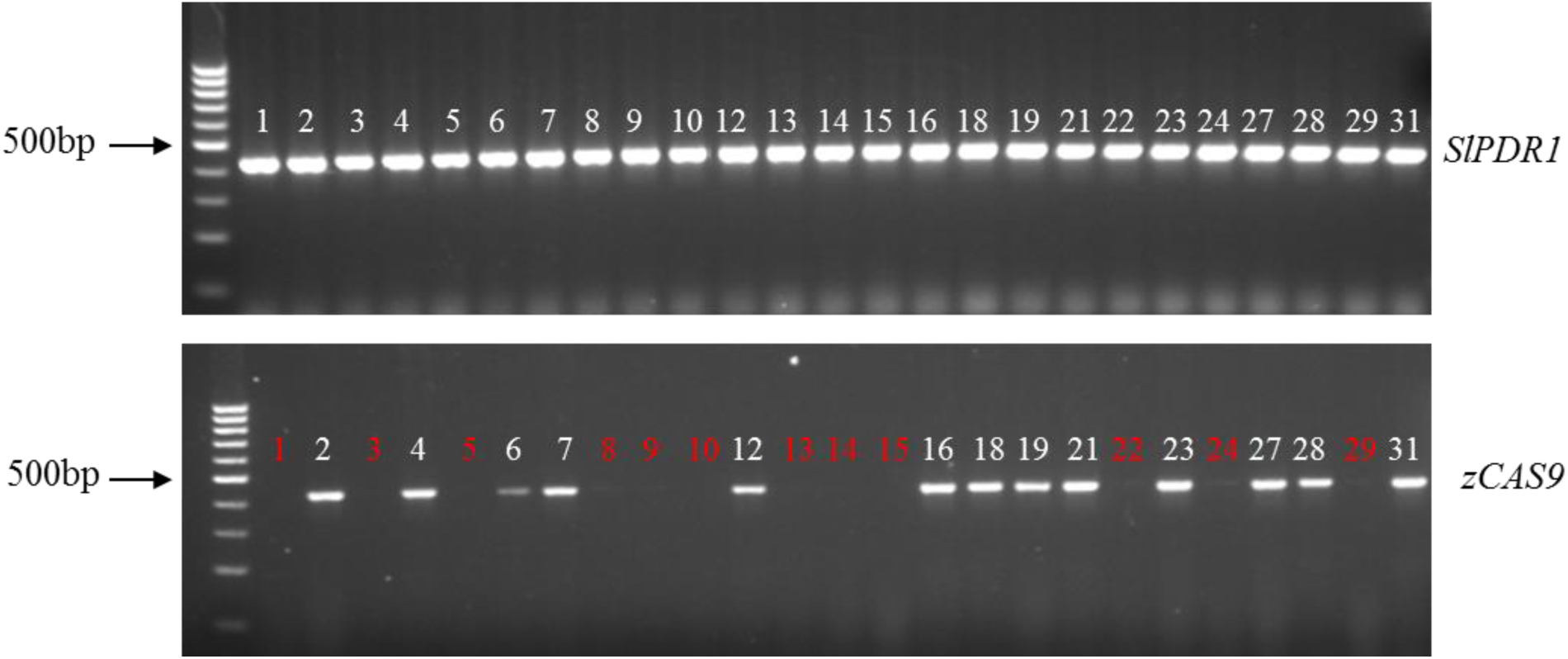
Identification of transgene free plants using zCas9 specific primer in T1 generation using different ^*ABC*^zCas9mutantsplants. Red color number denote transgene (zCas9) free plants.

**Supplementary Fig. S2.**
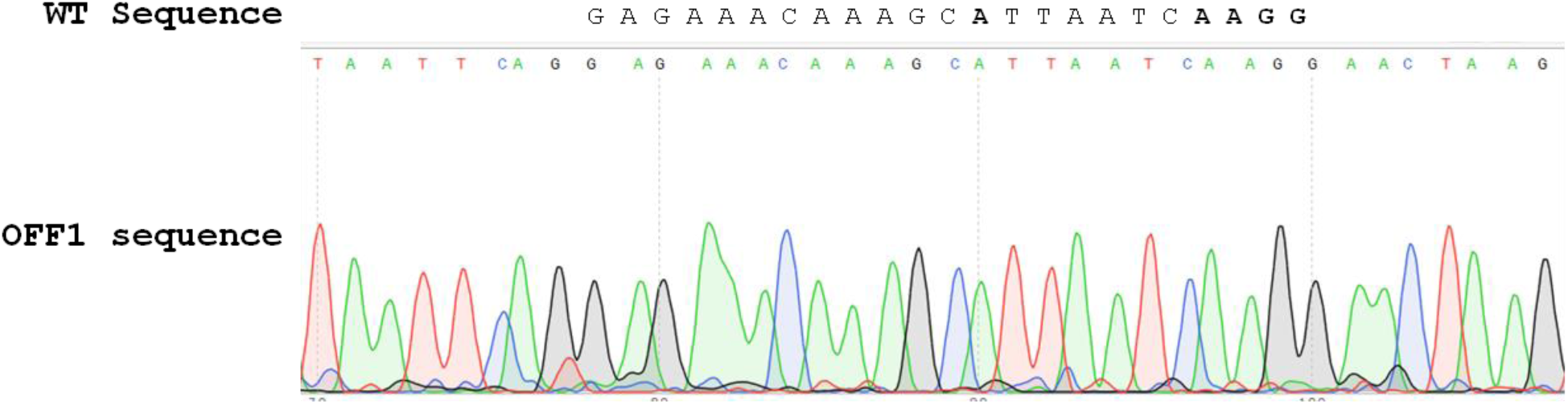
PCR product sequence chromatogram of potential off-target of the ABC-sgRNA OFF1 (*Solyc06g036240.2*). No off-target mutation detected in the genome of ABC-Cas9 edited lines. For sequencing PCR product, off target specific forward primers were used.

**Supplementary Fig. S3.**
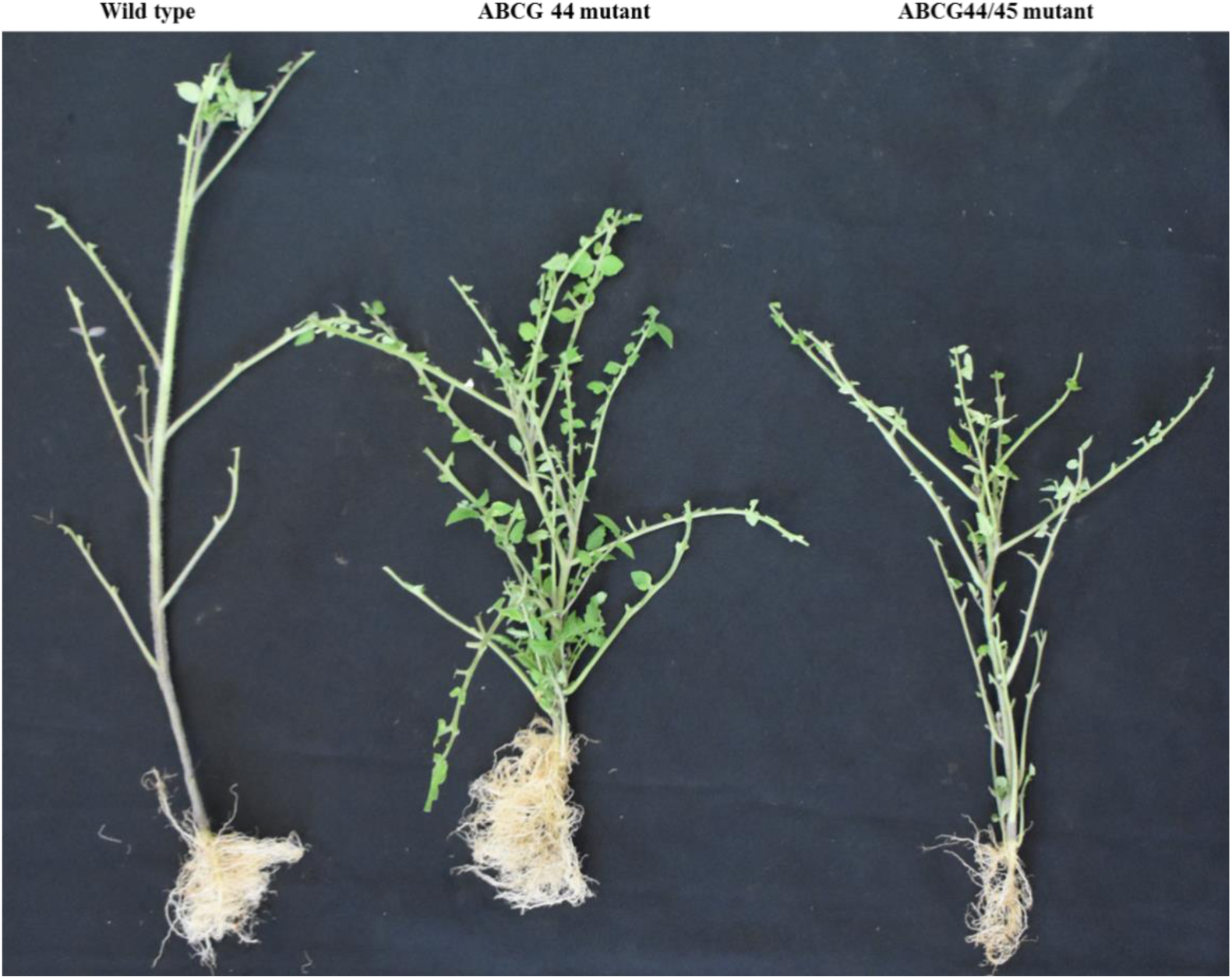
Characteristic shoot branching and axillary bud outgrowth phenotype of ^*ABC*^Cas9 knockout mutants in 1-month-old plants grown in green house.

**Supplementary Table S1.** Segregation pattern of ^*PDR1*^Cas9 mediated targeted mutation from T0 to the T1 generation. The zygosity of heterozygote in T0 plant lines were putative. +, Foreign DNA (zCas9) was detected; -, Foreign DNA (zCas9) was not detected.

| <sup>ABC</sup>sgRNA/Cas9<br>Locus | T0 generation |  | T1 generation |  |  |
| --- | --- | --- | --- | --- | --- |
|  | Mutation<br>type | Zygosity | Mutation<br>type/ total test | Mutation exist<br>no./ total test<br>(%) | Foreign DNA<br>(zCas9)<br>segregation<br>(%) |
| *Solyc08g067610* | -8nt/-4nt | Biallelic | 12(-8nt)<br>10 (Biallelic)<br>6(-4nt) | Homozygous<br>(18/28),<br>Biallelic<br>(10/28), (100%) | 13+; 12-<br>(48%) |
| *Solyc08g067620* | -1nt | Heterozygous | -1nt (<br>homozygous<br>5/25) | Homozygous<br>(5/25),<br>Heterozygous<br>(6/25), wt<br>(14/25) (50%) |  |

**Supplementary Table S2.**
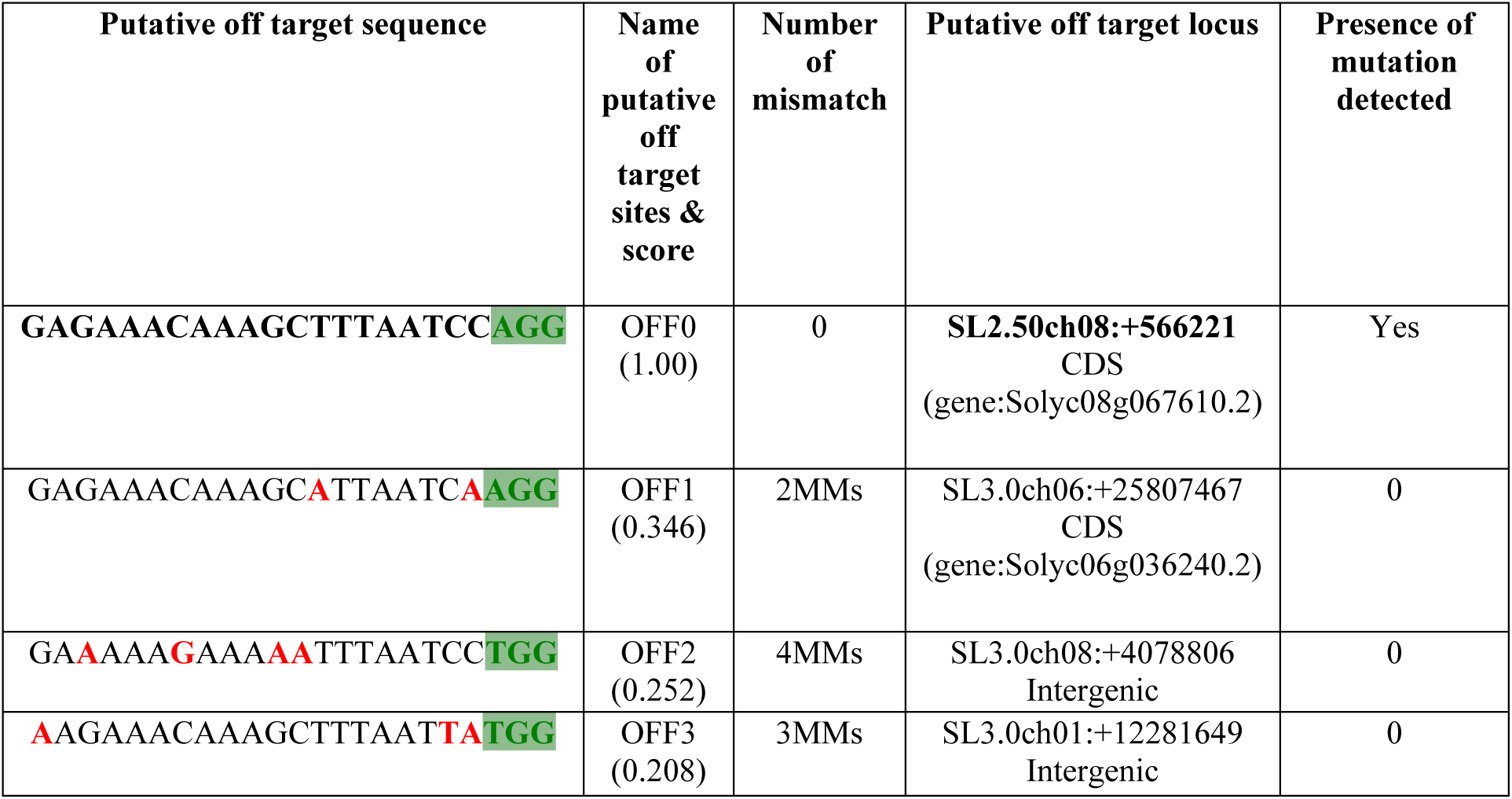
Mutation analysis of potential off-target sites in ABC mutated lines. To analyze the off target using CRISPR-P, high score value was considered as top hit, PAM sequence (NGG) is indicated in green, mismatch nucleotides are marked in red. Examined plants were randomly selected from the T1 generations of sgRNA-ABC/Cas9 edited tomato plants.

**Supplementary Table S3.**
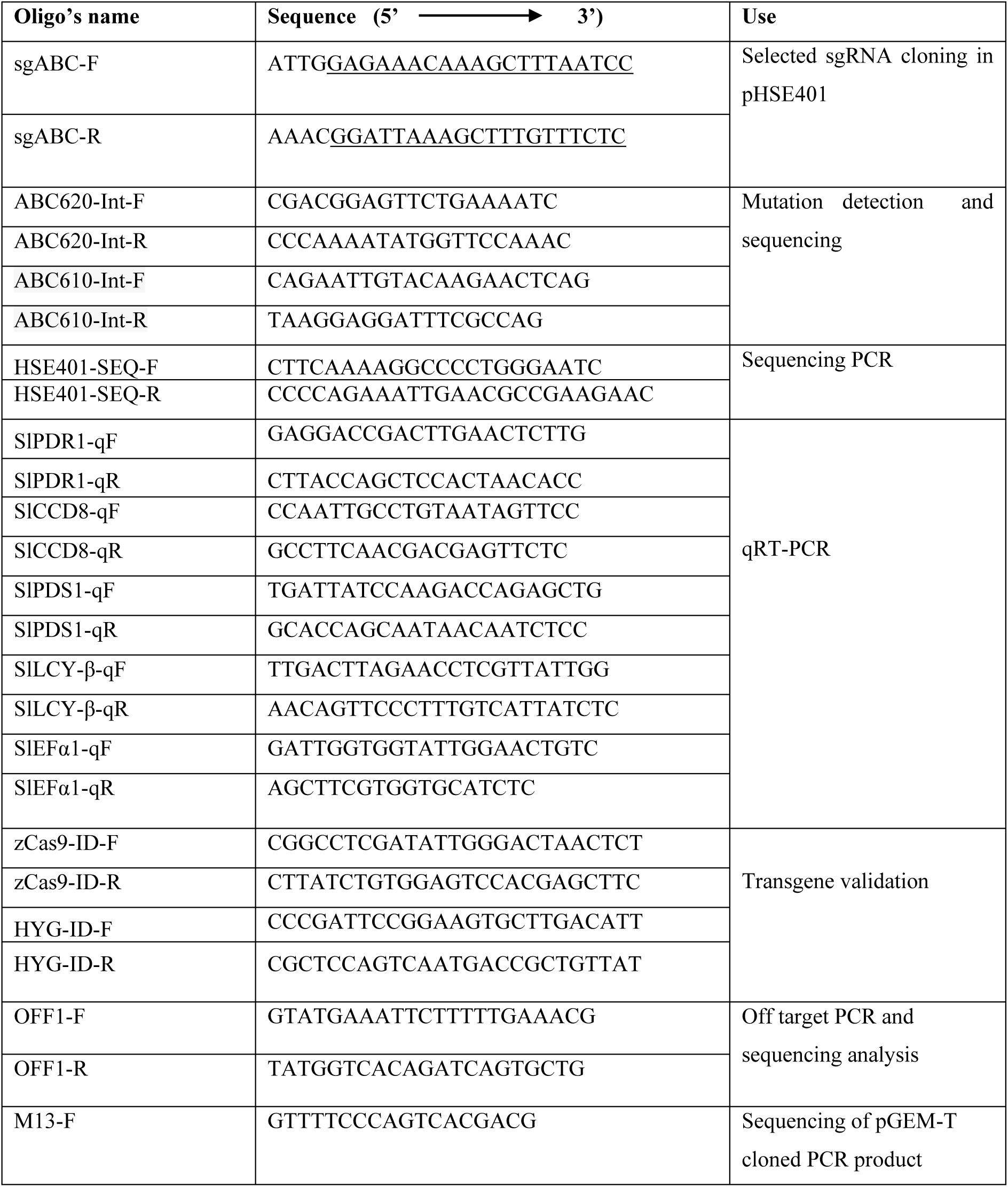
List of primers used in this study

